# Selecting stimulation intensity in repetitive transcranial magnetic stimulation studies: A systematic review between 1991 and 2020

**DOI:** 10.1101/2020.09.28.316190

**Authors:** Zsolt Turi, Maximilian Lenz, Walter Paulus, Matthias Mittner, Andreas Vlachos

**Author notes:** Shared senior authorship (Matthias Mittner, Andreas Vlachos).

## Abstract

**Background:** Repetitive transcranial magnetic stimulation (rTMS) is an increasingly used, non-invasive brain stimulation technique in neuroscience research and clinical practice with a broad spectrum of suggested applications. Among other parameters, the choice of stimulus intensity and intracranial electric field strength substantially impact rTMS outcome. This review provides a systematic overview of the intensity selection approaches and stimulation intensities used in human rTMS studies. We also examined whether studies report sufficient information to reproduce stimulus intensities in basic science research models. **Methods**. We performed a systematic review by focusing on original studies published between 1991 and 2020. We included conventional (e.g., 1 Hz or 10 Hz) and patterned protocols (e.g., continuous or intermittent theta burst stimulation). We identified 3,784 articles in total, and we manually processed a representative portion (20%) of randomly selected articles. **Results**. The majority of the analyzed studies (90% of entries) used the motor threshold (MT) approach and stimulation intensities from 80 to 120% of the MT. For continuous and intermittent theta burst stimulation, the most frequent stimulation intensity was 80% of the active MT. Most studies (92% of entries) did not report sufficient information to reproduce the stimulation intensity. Only a minority of studies (1.03% of entries) estimated the rTMS-induced electric field strengths. **Conclusion**. We formulate easy-to-follow recommendations to help scientists and clinicians report relevant information on stimulation intensity. Future standardized reporting guidelines may facilitate the use of basic science approaches aiming at better understanding the molecular, cellular, and neuronal mechanisms of rTMS.

## 1. Introduction

Repetitive transcranial magnetic stimulation (rTMS) non-invasively induces electromagnetic fields in the brain [1]. It is increasingly applied in neuroscience research and clinical treatment [2,3]. The efficacy of rTMS, besides the state of the receiving brain, significantly depends on the stimulation parameters, collectively referred to as the dose.

Based on the definition of Peterchev and colleagues [4], rTMS dose refers to all participant-independent device parameters that affect the spatial and temporal characteristics of the electromagnetic field produced in the brain. These include defining both the user-adjustable parameters (e.g., the percentage of maximum pulse intensity) and non-adjustable parameters (e.g., coil inductance) [4]. In this definition, the term “independent” refers to the notion that all stimulation parameters are translated into physical, participant-independent device parameters [4].

Selecting the stimulation intensity is an important step in determining rTMS dose. There are several intensity selection approaches, and researchers, as well as clinicians, may face two obvious challenges in this process. The first is conceptual: How to determine the stimulation intensity prospectively, such that the application of rTMS leads to the desired effects? The second is practical: For the sake of reproducibility, which parameters are crucial to standardize across experiments, and what information must be reported for a given protocol?

The goal of the present systematic review was to provide a comprehensive overview of the different intensity selection approaches. To this aim, we performed a careful literature search to identify the different approaches and quantify their relative frequency of use. Moreover, we evaluated whether studies report sufficient information required to reproduce the stimulation intensity. Reporting the stimulation parameters in a reproducible manner is essential for research and clinical applications [4]. Indeed, standardized reporting guidelines are crucial to informing computational and basic science research approaches to investigate the cellular, molecular, and network mechanisms of clinically relevant rTMS protocols [e.g., 5]. We formulate easy-to-follow recommendations to help improve the reporting quality of rTMS studies.

## 2. Methods

### 2.1. Creating the database

A systematic literature search was performed by identifying rTMS studies to create a comprehensive database. We searched for articles that were published between January 1, 1991, and July 31, 2020. To this aim, we used R (version 4.0.2) [6], RStudio integrated development environment (version 1.3.1073) [7], and the RISmed library (version 2.1.7) [8].

The following search terms were used: “repetitive transcranial magnetic stimulation,” “rTMS,” “repetitive TMS,” “rhythmic TMS,” “theta burst stimulation,” “TBS,” “cTBS,” and “iTBS.” The search revealed 6,727 PubMed hits in total, and we manually screened them to identify peer-reviewed original studies that applied rTMS on humans. We identified 3,804 articles that matched the criteria, and we could gain access to 3,784 articles. The remaining articles belonged to (i) other article types (e.g., reviews), (ii) animal studies, (iii) in-vitro studies, or (iv) did not use rTMS.

### 2.2. Randomly sampling from the database

Random samples were taken from the accessed articles using a multi-step procedure (for an overview, see Table S1). In each step, we sampled and processed 380 articles, which corresponds to ~10% of the total number of articles. After the second step, we compared the result with the combined results of the first and second steps. We assessed similarity and compared the results from the steps based on the stimulation intensity selection approaches. If the results were not different, we interpreted the findings to indicate that the results could be generalized to the entire database. In case of a discrepancy, we planned to continue with the next step until we observed no difference in the results between the last and second-last steps. However, this was not necessary, because we observed highly similar results between the first and second sampling (see Fig. S1). Thus, we processed 760 articles in total.

### 2.3. Extracting the stimulation parameters

We performed a text-mining search to identify the stimulation intensity approaches, the stimulation intensity, and device-specific information using R library pdfsearch (version 0.3.0) [9] on the 760 randomly selected articles. Moreover, we also evaluated on the entire database, whether studies report the electric field values (see Table S2 for details). Note that we only used the text-mining search to facilitate the manual information extraction process; all information for each study was manually assessed.

Only real stimulation protocols were considered. Consequently, sham/control protocols were not included in this review. Because the present review focused on how rTMS studies determine the stimulation intensity and its relationship to the stimulation frequency, we created a new entry for every unique combination of the stimulation frequency and intensity in a given study. For instance, we created two separate entries for the scenario, when a given study used 1 Hz rTMS at 90 and 110% of the resting motor threshold.

However, studies frequently manipulated other stimulation parameters, such as anatomical target or pulse number. We added each stimulation parameter as a separate entry up to three values; that is, up to three anatomical targets. For example, we created two separate entries for the scenario, when 1 Hz rTMS at 90% of the resting motor threshold was applied over the prefrontal and parietal cortices. Beyond this value (i.e., three anatomical targets), we treated the protocol as a single entry and labeled it using more than three values for a given parameter. This condition is commonly applied for mapping studies, such as pre-operative mapping of eloquent brain areas.

## 3. Results

### 3.1. Intensity selection approaches

The number of original studies steadily increased during the observation period, reaching 250–300 publications per year during the past 5 years (see Figure 1). We processed 760 randomly sampled articles, identified 241 unique protocols and added 1,378 entries in total. Because the details of the stimulation intensity parameters may differ between the entries in a given study, we decided to report the number of entries, not studies, unless stated otherwise. Consequently, the associated percentage values refer to the percentage of the total number of identified entries.

**Figure 1.**
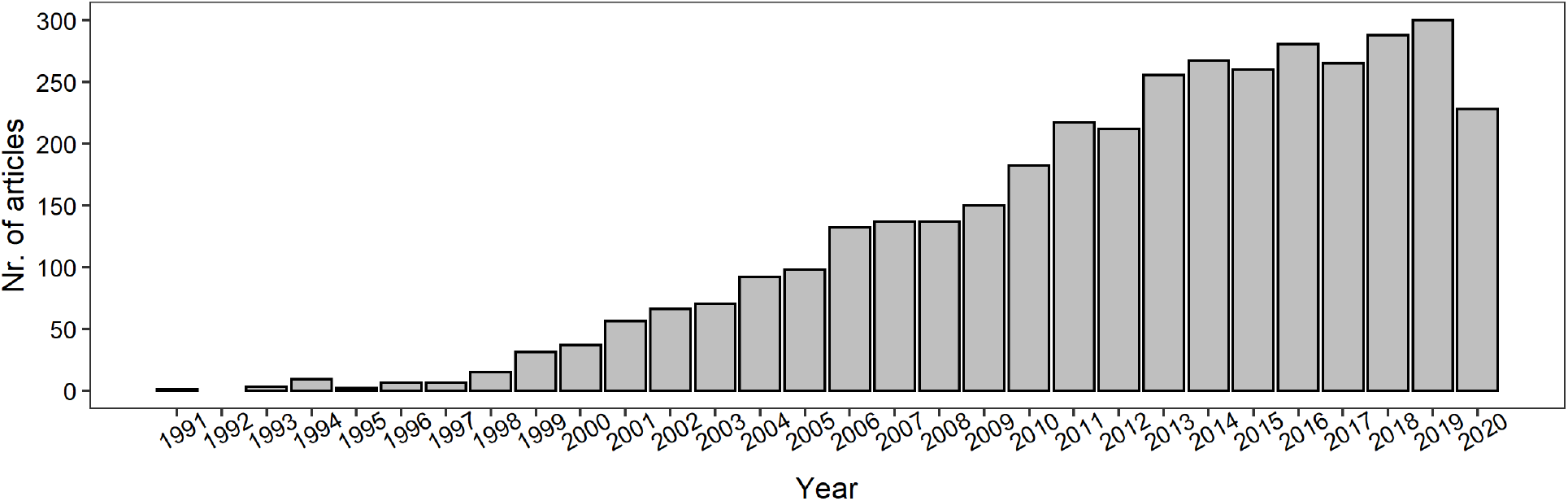
Number of identified original studies performing rTMS on humans. The number of articles per year is shown between January 1, 1991 and July 31, 2020.

We identified three main stimulation intensity selection approaches: 1) threshold based (n = 1,257; 91.22%), 2) fixed intensity (n = 91; 6.6%), and 3) electric field estimation based (n = 2; 0.15%). A small portion of entries failed to report the approach (n = 28; 2.03%). For an overview, see Figure 2.

**Figure 2.**
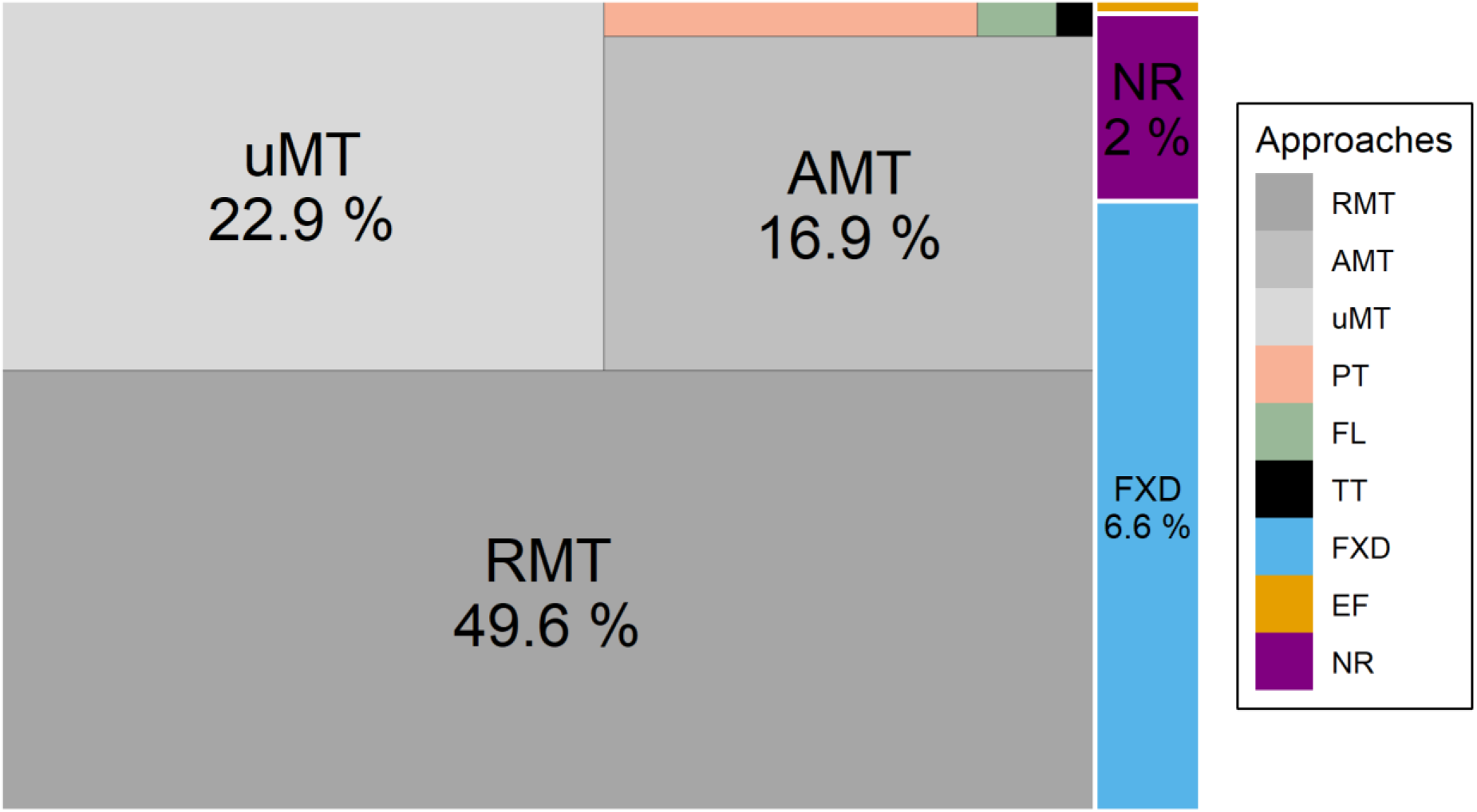
Stimulation intensity selection approaches and their relative frequency of use, by percentage of the total number of identified entries (n = 1,378). The sizes of rectangles are proportional to the percentage values. Abbreviations: RMT: AMT: active motor threshold, resting motor threshold, uMT: unspecified motor threshold, PT: phosphene threshold, FL: functional lesion threshold, TT: tolerability threshold, FXD: fixed intensity, EF: electric-field-based intensity selection, NR: not reported.

The most common threshold-based approach was the motor threshold (MT) approach (n = 1,232; 89.40%). Most studies used the resting threshold (n = 683; 49.56%) or the active MT approach (n = 233; 16.91%). Some studies did not specify the type of threshold (n = 316; 22.93%). Throughout the article, we use the abbreviation MT to collectively refer to the active, resting, and unspecified motor threshold. Otherwise, we use the adjectives active, resting, or unspecified to explicitly refer to a given subtype of the MT.

Most studies used the method of limit (n = 776, 56.31%) to estimate the MT. In this approach, the minimal device output is determined by decreasing or increasing the device output required to produce motor evoked potentials with a minimum predefined peak-to-peak amplitude 50% of the time in a given number of trials [10]. Some studies did not specify the exact method; instead, they referred to previous publications via citation (n = 107, 7.76%). A small subset of studies used other algorithms to estimate the MT (e.g., parameter estimation by sequential testing methods) [10,11] (n = 28, 2.03%). Many studies did not report any details of the method used to estimate the MT (n = 340, 24.67%).

In some studies, the MT was determined by recording motor evoked potentials with surface electrodes (n = 705, 51.16%). In other studies, MT was visually identified by observing the muscle twitches (n = 177, 12.84%). In many cases, the authors provided no information about how the MT was detected (n = 350, 25.40%).

Some studies used additional, although less common, threshold-based approaches. A small number of studies used the phosphene threshold (n = 19, 1.38%), the threshold for inducing “functional lesions” (n = 4, 0.29%), such as speech arrest, or the tolerability threshold (i.e., the maximum intensity of comfortable stimulation; n = 2, 0.15%) to set the stimulation intensity.

### 3.2. Most studies use the motor threshold approach at 80 to 120% of the threshold

The most frequent intensity selection approach was the MT-based one. Here, the stimulation intensity is typically expressed as the percentage (e.g., 90%) of the MT. Two methods are commonly used for estimating the MT. The active MT requires the slight contraction of the target muscle by ~10–20% of the maximum contraction intensity. The resting MT is assessed during complete relaxation of the target muscle.

We selected articles that (i) used the MT approach, (ii) used a single stimulation intensity, and (iii) reported the stimulation intensity and frequency (n = 1,162, 84.33%). Of these studies, the stimulation intensity typically ranged from 80 to 120% of the MT (Figure 3). Studies using resting MT were the most frequent (Figure 3B).

**Figure 3.**
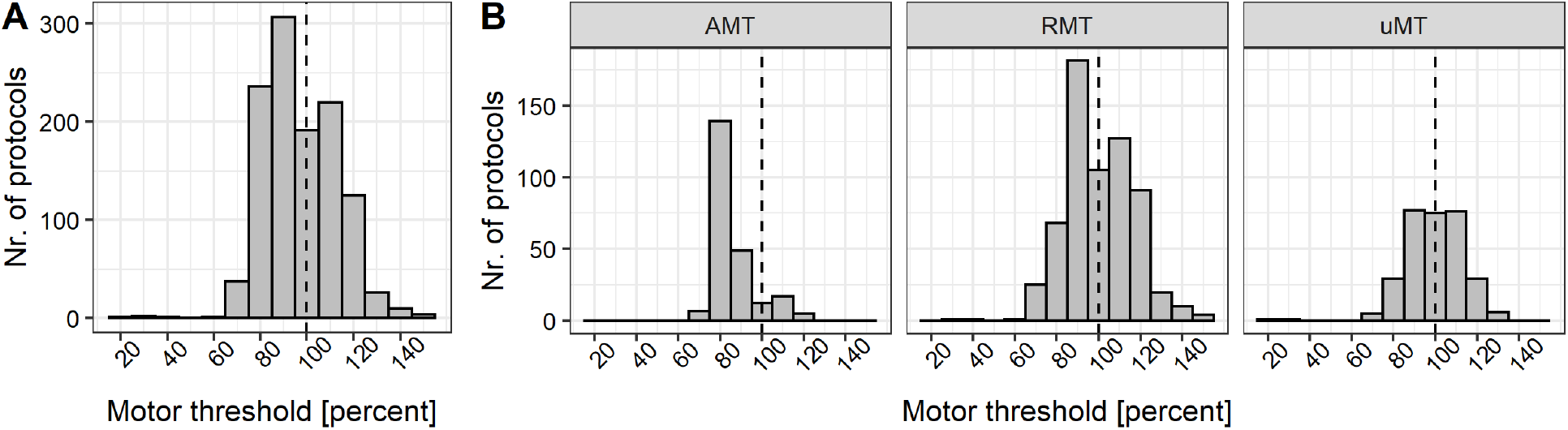
Stimulation intensity selection based on the motor threshold approach. Motor threshold percentages typically ranged from 80 to 120%. Vertical dashed line highlights the intensity corresponding to the 100% motor threshold. Abbreviations: AMT: active motor threshold, RMT: resting motor threshold, uMT: unspecified motor threshold.

### 3.3. Stimulation intensity as a function of stimulation frequency

Next, we focused on the relationship between the stimulation intensity and stimulation frequency. We used the same selection criteria described in the previous point.

We divided the rTMS protocols into conventional and patterned ones. Conventional protocols use single frequencies (e.g., 1 Hz) [12], whereas patterned protocols combine at least two stimulation frequencies (e.g., 5 and 50 Hz as in theta burst rTMS) [13]. Moreover, patterned protocols use standardized stimulation parameters, such as inter burst intervals. We also identified studies using quadripulse protocols that deliver four monophasic pulses with interpulse intervals of 5 or 50 ms [14,15]. However, due to the low occurrence rate in our sample (n = 5, 0.36%), we did not analyze these protocols in detail.

Focusing on the conventional protocols, we found that during the past 20 years, 1 Hz stimulation protocols were most frequently used (n = 304, 22.06%), followed by 10 Hz (n = 230, 16.70%), 5 Hz (n = 126, 9.14%), 20 Hz (n = 122, 8.85%), 25 Hz (n = 22, 1.60%), and 15 Hz (n = 17, 1.23%). See also Figure 4.

**Figure 4.**
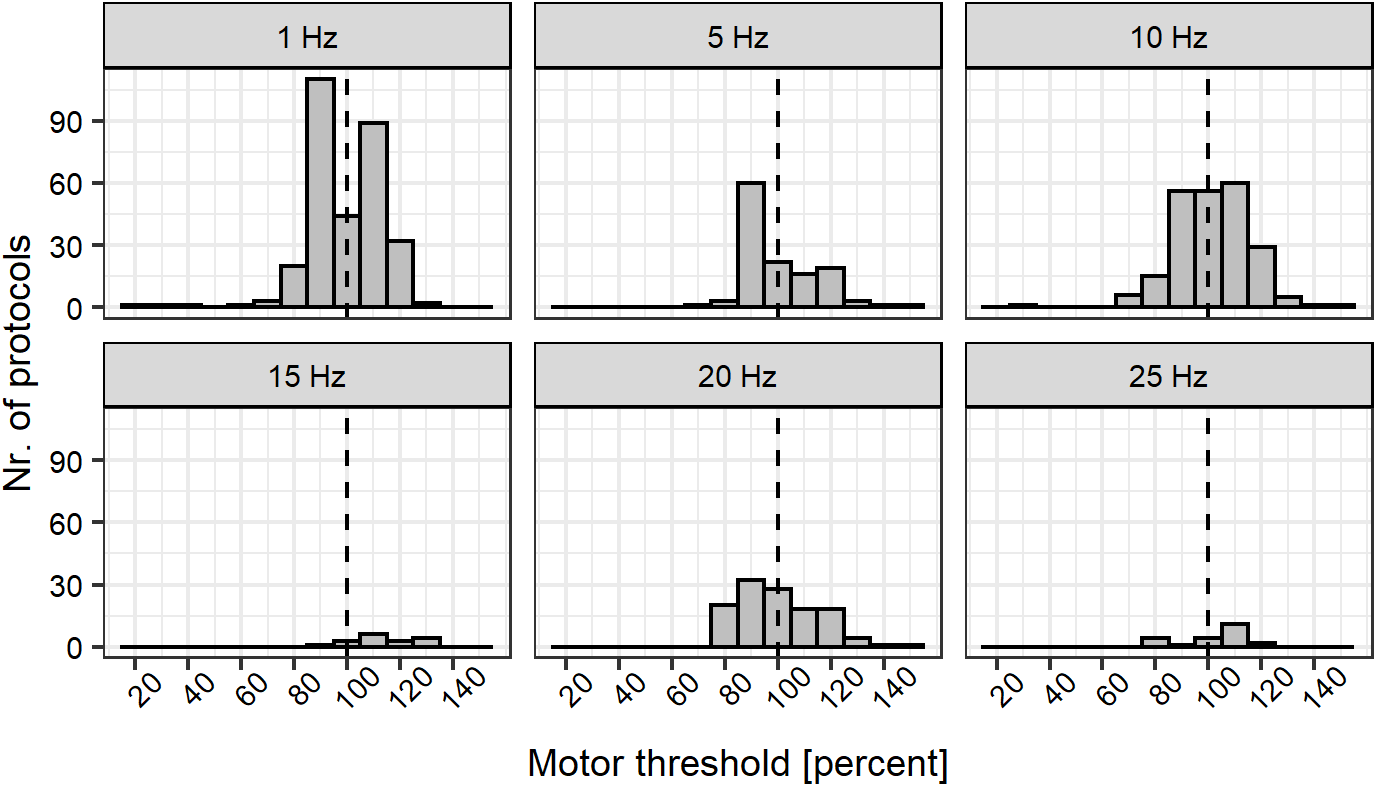
Stimulation intensity expressed as a percentage of motor threshold in the conventional protocols arranged according to stimulation frequency. Note the range of intensities and differences between protocols. Intensities correspond to the resting, active, and unspecified motor thresholds.

Although the stimulation intensity typically ranged from 80 to 120% of the MT, we found “popular” stimulation intensities for specific stimulation frequencies, shown in Figure 4. The substantial variability in the stimulation intensity selection approaches makes the direct comparison between the various protocols difficult. For example, the most common stimulation intensities were 90 and 110% of the MT for 1 Hz protocols. For the 5 Hz protocol, the most frequent intensity was 90% of the MT. For 10 Hz, the most frequent intensities ranged from 90 to 110% of the MT. Note that the 10 Hz protocol approved by the U.S. Food and Drug Administration (FDA) uses 120% of the resting MT [16].

We found that the theta burst protocols (n = 210, 15.24%) used lower stimulation intensities than conventional protocols. The most common stimulation intensity was 80% of the active MT (Figure 5). Whereas for the conventional protocol, the resting MT was the most frequent intensity selection approach, for the theta burst protocols, the active MT was the dominant approach (Figure 6). Note that the U.S. FDA-approved intermittent theta burst protocol uses 120% of the resting MT [3].

**Figure 5.**
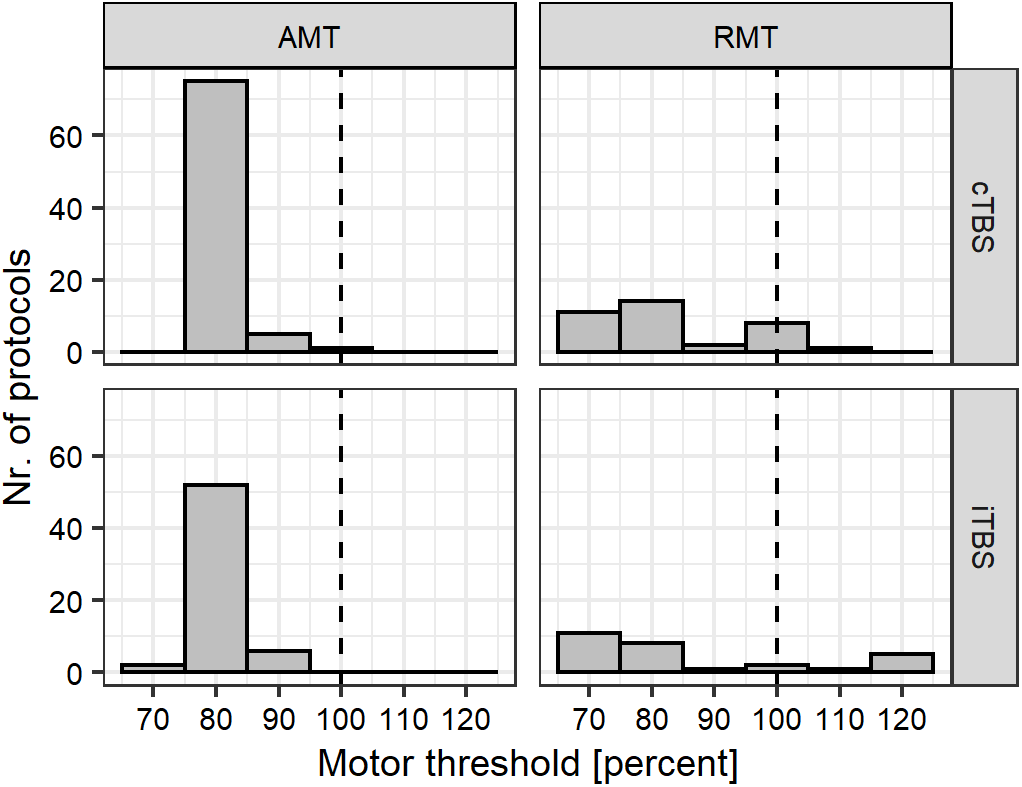
Stimulation intensity expressed as percentages of motor threshold assessed in theta burst protocols. Abbreviations: AMT: active motor threshold, RMT: resting motor threshold, cTBS: continuous theta burst stimulation, iTBS: intermittent theta burst stimulation.

**Figure 6.**
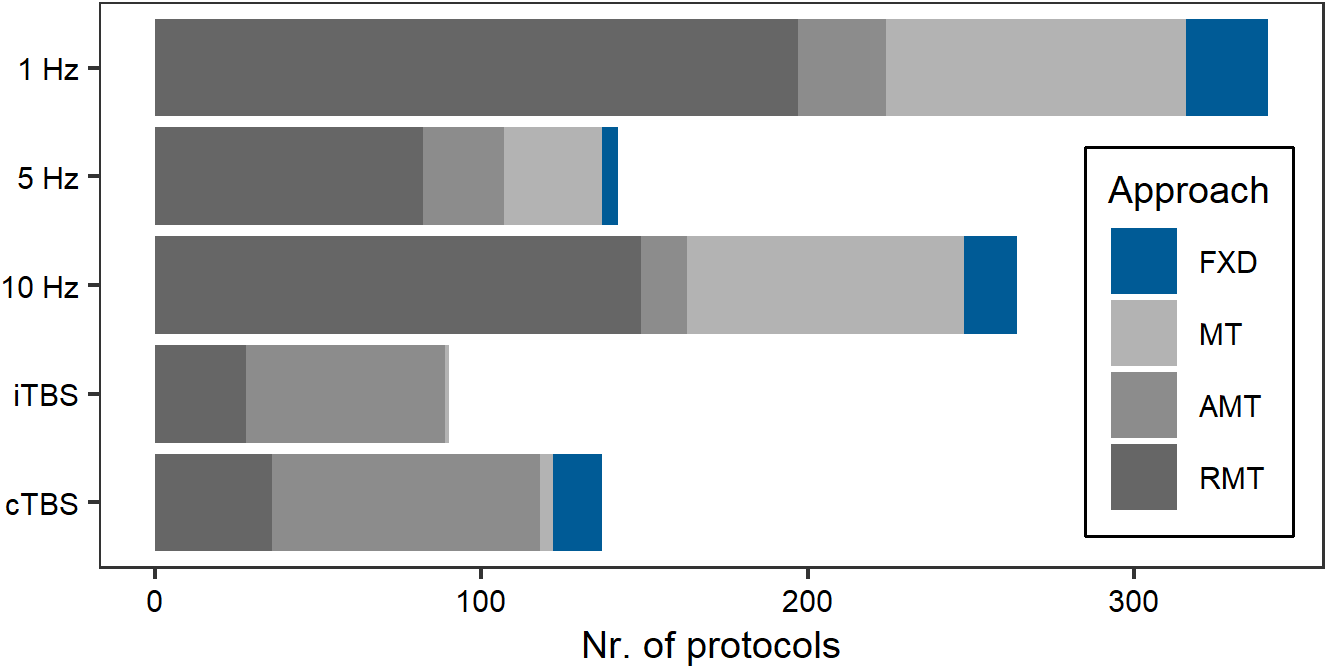
Relative frequency of motor threshold intensity selection approaches in conventional (1, 5, and 10 Hz) and theta burst protocols. Abbreviations: AMT: active motor threshold, RMT: resting motor threshold, uMT: unspecified motor threshold, FXD: fixed stimulation intensity, cTBS: continuous theta burst stimulation, iTBS: intermittent theta burst stimulation.

### 3.4. Reproducibility of stimulation intensity

The seminal work of Peterchev and colleagues in 2012 [4] emphasized the importance of reporting all user-adjustable and device-specific information that influences the resulting electromagnetic field. Here, we focus on the stimulation intensity from the perspective of basic science and electric field estimation approaches. We acknowledge that reproducing the dose (and not only the stimulation intensity) is a more complex and challenging task beyond the scope of the present review.

In basic science and computational approaches, the goal is to fine-tune the stimulation parameters in a manner that closely matches the produced electromagnetic fields utilized in clinically relevant rTMS protocols. To ensure the reproducibility of the stimulation intensity, it is necessary to report (i) the individual percentages of the maximum stimulator output and (ii) the company and model identification number of the rTMS device. Furthermore, in human studies, one should also describe the (iii) intensity selection approach, (iv) its detailed procedure, and (v) the stimulation intensity (e.g., 90% of the resting MT). If applicable, human studies should share the defaced, anonymized, anatomical magnetic resonance imaging (MRI) data of the participants.

We determined how many studies reported all information required to reproduce the stimulation intensity (points i–ii). We identified only 110 entries (7.98%) that fulfilled these criteria.

We found that only a highly limited number of entries reported all the criteria described above (points i-v; n = 57, 4.14%). Stricktly speaking, the reported stimulation protocol is not reproducible if any of the abovementioned parameters (i.e., points i-v) are missing from the description. Strikingly, in more than half of the entries assessed here, the device output or coil model was not reported. This information is essential for approximating the induced electric fields in a given head model [1]. Table 1 summarizes the missing information and its frequency.

**Table 1.**
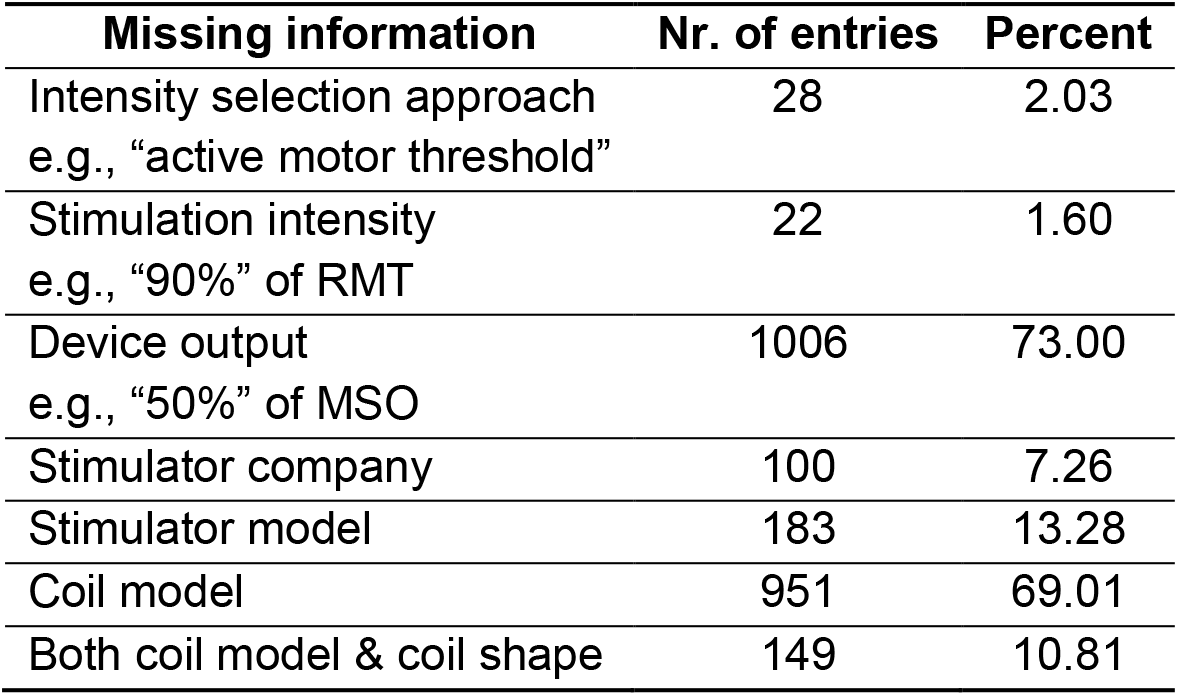
Missing information required to reproduce rTMS stimulation intensity.

### 3.5. Reporting electric field strength

We identified only 39 out of 3,784 studies (1.03%) that reported the electric field strength in the brain produced by rTMS. The reported electric field values ranged from 14 to 182 V/m. Most studies reported the estimated electric field given by the neuronavigation system (n = 31, 0.82%). Six used the finite element method (simulation of non-invasive brain stimulation - SimNIBS, openly available since 2015 [17]), one used the boundary element method, and one a spherical head model. Most studies (n = 36, 0.95%) set the stimulation intensity based on the MT and then estimated the corresponding electric field strength. We found only three studies that determined the stimulation intensity based on estimating the electric field strength.

Notably, more studies reporting the electric field strength have been published within the past 10 years, indicating an increased requirement for computational tools to estimate and standardize electric fields induced by a given rTMS.

## 4. Discussion

In this review, we focused on how human rTMS studies determine the stimulation intensity for rTMS. To this aim, we created a database comprising 3,784 original studies and manually processed 760 randomly selected articles.

We found that the most common intensity selection approach was the motor threshold one. Most studies used a single stimulation intensity (e.g., 110% of the resting motor threshold). In contrast, a minority of studies used a range of stimulation intensities (e.g., 100–110% of the resting motor threshold).

The 1, 10, and 5 Hz were the three most popular conventional protocols. We found that most protocols used stimulation intensities from 80 to 120% of the motor threshold. Theta burst protocols used slightly lower stimulation intensities, typically at 80% of the active motor threshold. Whereas the threshold-based estimation approaches used individualized stimulation intensities, the less common fixed-intensity approach delivered a predetermined stimulation intensity (e.g., 50% of the device output).

According to the current reporting convention, the vast majority of articles (ca. 92%) did not report sufficient information required to readily reproduce the stimulation intensity and approximate the electric field for rTMS.

### 4.1. Need to link dose to electric field

Several factors can contribute to selecting the stimulation intensity in human rTMS studies. The most frequent factors are (i) conventions of the field (e.g., using the resting motor threshold even when targeting a different cortical region than the motor cortex itself), (ii) the safety and tolerability of the participants, and (iii) device limitations. Surprisingly, the electric field rarely played an explicit role in determining the stimulation intensity for rTMS [but see 18–20].

The variability in the electric field strength of non-invasive brain stimulation techniques significantly affects the underlying neural mechanisms [21]. Electric field simulations play an essential role in linking *in vitro* and *in vivo* studies targeting cell cultures, brain slices, or the entire, *in situ* brains of different species [5,22]. The lack of reporting the electric field strength and participant-independent device parameters in human rTMS studies hinders the translational use of computational and basic science approaches. Mutli scale neuronal modeling is crucial in informing studies on the molecular, cellular, and neural mechanisms of rTMS (e.g., [23]).

For instance, estimating the resting motor threshold in rodents is different from the procedure typically used in humans due to anesthetics [c.f., 24]. Similarly, it is uninformative when *ex vivo* studies such as brain slice experiments define the stimulation intensity exclusively using percentages of the motor threshold [5]. In this context, reporting the corresponding device output for the stimulation intensity and the electric field is of utmost importance. Translating the motor threshold in human participants to *in vitro* basic research settings is hardly possible.

However, there is substantial variability in how one can describe the resulting electric field properties. Because the produced electric field is a complex three-dimensional vector field, it is not possible to characterize it using a single parameter [4]. Furthermore, there is an ongoing debate about which component of the rTMS-induced electric field produces the relevant physiological effects [e.g., 25,26].

Based on these considerations, it appears to be important to share all necessary information and data required to reproduce not only the dose but also the electromagnetic field induced by a given rTMS protocol [e.g., 18]. As the accuracy and user-friendliness of electric field simulations improve, we expect an increase in the number of studies that standardize certain electric field properties across subjects.

### 4.2. Recommendations for reproducibility

Our analyses revealed that the overwhelming majority of studies (92%) did not report sufficient information to reproduce the stimulation intensity. Our findings agree with the commentary of Wilson and St. George [27], who draw attention to the variability and lack of methodlolgical transparency and standardization in estimating and reporting motor evoked potentials in human studies.

One way to overcome the limitations of the current reporting convention is to establish a consensus about standardized reporting guidelines. Editors of specialized peer-reviewed journals, who are often the leading experts in the field, can play an important role in developing and enforcing such guidelines.

At the same time, companies selling the devices should ensure access to all device-specific information (e.g., coil inductance). The model identification number should be readily available for the user to facilitate easy referencing. Thus, the authors could only report the user-adjustable parameters while referring to non-adjustable, fixed-device parameters via unique device identification numbers. The goal of such a referencing system is that authors do not have to deal with a long list of nuanced but important device-specific parameters.

Below, we list our recommendations from a basic science perspective, which may serve as an orientation for future initiatives aiming to define the relevant standards in the field (see Figure 7). We focused on the motor threshold approach because it is unequivocally the dominant approach in the literature.

**Figure 7.**
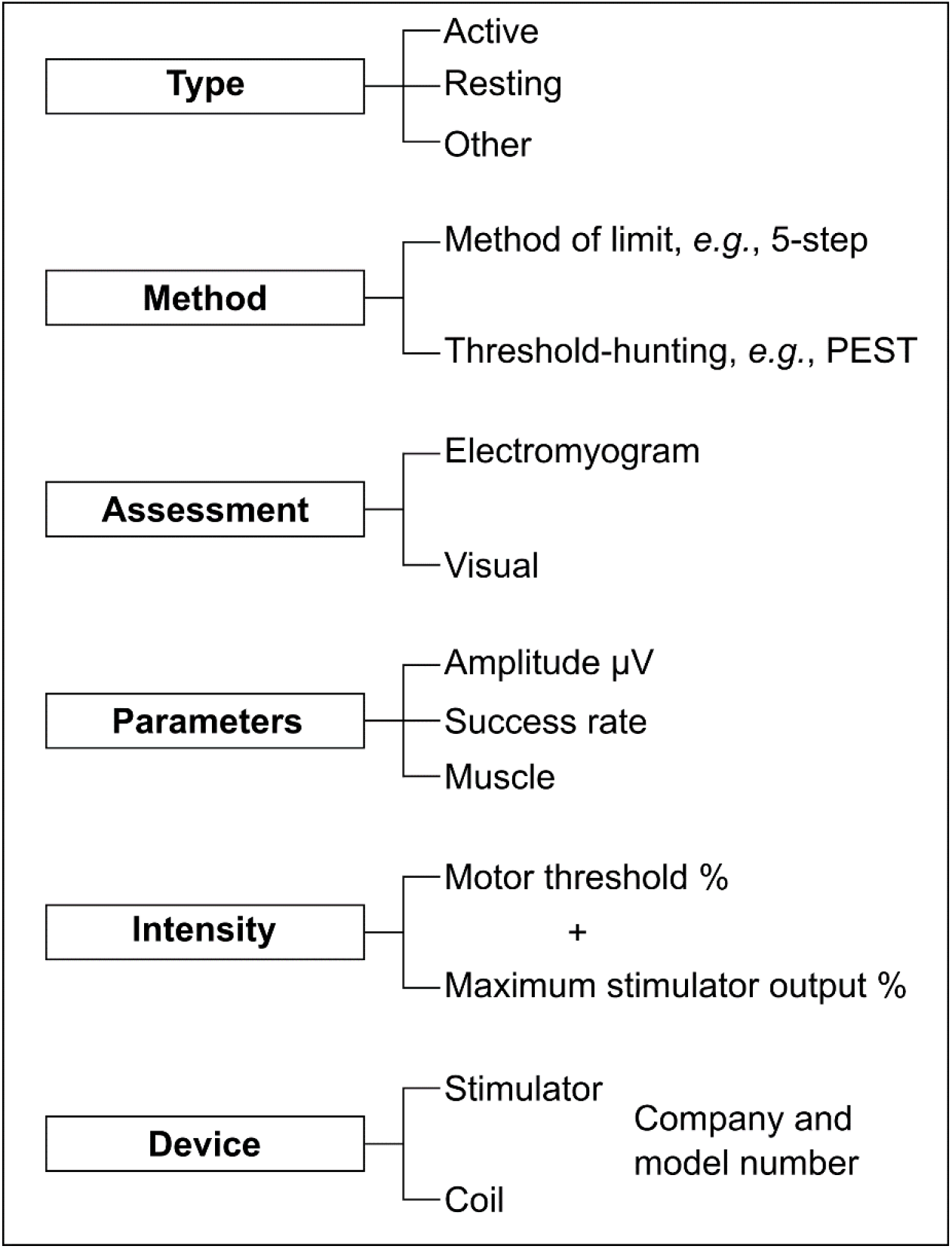
Crucial information to report when using the motor threshold intensity selection approach.

#### Recommendation 1

Many studies reported using the motor threshold approach without specifying the type of motor threshold and the exact methods of how it was determined. Different motor threshold approaches yield varying stimulation intensities [28] and electric field strengths. Therefore, it is essential to report explicitly about whether the motor threshold was an active, resting motor threshold or based on another approach, (e.g., to produce interhemispheric inhibition) [29].

#### Recommendation 2

One can estimate the threshold using various methods, such as the method of limit or threshold hunting algorithms [11,30,31]. Many studies do not mention the method at all; instead, they only state that the motor threshold “was established according to published criteria.” Alternatively, the authors only name the method (e.g., method of limit) and refer to the procedure via citation. One obvious limitation of this reporting convention is that the article may not be available at all research institutes. Because word limitations are typically not imposed on the methods section, we recommend explicitly reporting the method’s details.

#### Recommendation 3

Many studies used the method of limit to estimate the active or resting motor threshold. Although many studies used an electromyogram to record the amplitude of the motor-evoked potentials, the exact values often remained undefined. For example, the minimum amplitude for the active motor threshold may be between 100 and 200 μV. A typical minimum value for the resting motor threshold is 50 μV.

Another important aspect is the success rate because the exact number of successfully evoked motor potentials above a certain amplitude may vary. The typical values are 3 of 5 or 5 of 10 TMS pulses. However, we observed many other success rates. Success rates were often not reported. Therefore, we recommend reporting the peak-to-peak threshold amplitude and the success rate when using the method of limit.

#### Recommendation 4

Most studies used the motor threshold intensity selection method. Studies usually reported the stimulation intensity as a certain percentage of the motor threshold (e.g., 90% of the resting motor threshold). Although describing the method of selecting the stimulation intensity is an important step, reporting the associated stimulation intensities in terms of the device output is crucial for basic science and electric field simulation approaches. For example, the active motor threshold may correspond to 36% of the device output in one participant and 59% in another [32]. Therefore, it is essential to report the device output, typically referred to as the maximum stimulator output percentage.

#### Recommendation 5

It is a common shortcoming that studies only state the manufacturer of the stimulator and the shape of the coil (e.g., figure-of-eight-shaped coil). However, the stimulator and coil parameters (e.g., coil inductance) can significantly affect the produced electromagnetic field. Different coils attached to the same stimulator may end up in different results [33]. By accurately reporting the exact stimulator and coil model, one can trace these important parameters.

### 4.3. Conclusion

Due to the extensive set of possible parameters, it is currently challenging to decide which parameters to report for rTMS. Consequently, the overwhelming majority of studies do not report sufficient information to reproduce the stimulation intensity and, eventually, the dose of rTMS. This failure to report stimulation parameters makes it nearly impossible to properly reproduce published protocols and design appropriate basic science experiments. There is a need for developing a standardized and easy-to-follow reporting guideline to document the most crucial stimulation parameters in the best possible way.

## 5. Acknowledgment

We thank Swantje Reifarth and Alina Seiler from the Department of Clinical Neurophysiology, University Medical Center Göttingen, Georg-August University, Göttingen, Germany, for their assistance in initially screening the articles. We thank Celina Jacobsen and Sarjo Kuyateh from the Institute of Psychology, UiT The Arctic University of Norway, for their assistance in manually processing the articles. We thank Lisa Kuhlmann and Anna Wolfers from the Department of Neuroanatomy, Institute of Anatomy and Cell Biology, University of Freiburg, Germany, for their assistance in manually processing the articles. The study was, in part, supported by the University Medical Center Göttingen starting grant (to ZT) and an NIH-RO1 grant (1R01NS109498-01A1 to AV).

## 6. Authors contribution

Authors contribution was prepared according to the Contributor Roles Taxonomy. Conceptualization: ZT; Formal analysis: ZT; Funding acquisition: ZT and AV; Investigation: ZT; Methodology: ZT, MM and AV; Project administration: ZT, WP, MM and AV; Software: ZT; Supervision: ZT and AV; Visualization: ZT; Writing - original draft: ZT, ML, WP, MM and AV.

## Notes

### Competing Interest Statement

The authors have declared no competing interest.

## References

[1] Saturnino GB, Puonti O, Nielsen JD, Antonenko D, Madsen KH, Thielscher A. SimNIBS 2.1: A Comprehensive Pipeline for Individualized Electric Field Modelling for Transcranial Brain Stimulation. Cham (CH): Springer; 2019. https://doi.org/10.1007/978-3-030-21293-3.

[2] Suppa A, Huang YZ, Funke K, Ridding MC, Cheeran B, Di Lazzaro V, et al. Ten Years of Theta Burst Stimulation in Humans: Established Knowledge, Unknowns and Prospects. Brain Stimul 2016;9:323–35. https://doi.org/10.1016/j.brs.2016.01.006.

[3] Blumberger DM, Vila-Rodriguez F, Thorpe KE, Feffer K, Noda Y, Giacobbe P, et al. Effectiveness of theta burst versus high-frequency repetitive transcranial magnetic stimulation in patients with depression ( THREE-D): a randomised non-inferiority trial. Lancet 2018;391:1683–92. https://doi.org/10.1016/S0140-6736(18)30295-2.

[4] Peterchev AV, Wagner TA, Miranda PC, Nitsche MA, Paulus W, Lisanby SH, et al. Fundamentals of transcranial electric and magnetic stimulation dose: Definition, selection, and reporting practices. Brain Stimul 2012;5:435–53. https://doi.org/10.1016/j.brs.2011.10.001.

[5] Lenz M, Galanis C, Mu F, Opitz A, Wierenga CJ. Repetitive magnetic stimulation induces plasticity of inhibitory synapses. Nat Commun 2016:7:10020. https://doi.org/10.1038/ncomms10020.

[6] R Core Team. R: A Language and Environment for Statistical Computing 2020.

[7] RStudio Team. RStudio: Integrated Development Environment for R 2020.

[8] Kovalchik S. RISmed: Download Content from NCBI Databases 2017.

[9] LeBeau B. pdfsearch : Search Tools for PDF Files. J Open Source Softw 2018;2:1–2. https://doi.org/10.21105/joss.00668.

[10] Borckardt JJ, Nahas Z, Koola J, George MS. Estimating Resting Motor Thresholds in Transcranial Magnetic Stimulation Research and Practice. J ECT 2006;22:169–75. https://doi.org/10.1097/01.yct.0000235923.52741.72.

[11] Awiszus F. Using relative frequency estimation of transcranial magnetic stimulation motor threshold does not allow to draw any conclusions about true threshold. Clin Neurophysiol 2014;125:1285–7. https://doi.org/10.1016/j.clinph.2013.09.046.

[12] Chen R, Classen J, Gerloff C, Celnik P, Wassermann EM, Hallett M, et al. Depression of motor cortex excitability by low-frequency transcranial magnetic stimulation. Neurology 1997;48:1398–403. https://doi.org/10.1212/WNL.48.5.1398.

[13] Huang Y-Z, Edwards MJ, Rounis E, Bhatia KP, Rothwell JC. Theta burst stimulation of the human motor cortex. Neuron 2005;45:201–6. https://doi.org/10.1016/j.neuron.2004.12.033.

[14] Hamada M, Hanajima R, Terao Y, Arai N, Furubayashi T, Inomata-Terada S, et al. Quadro-pulse stimulation is more effective than paired-pulse stimulation for plasticity induction of the human motor cortex. Clin Neurophysiol 2007;118:2672–82. https://doi.org/10.1016/j.clinph.2007.09.062.

[15] Matsumoto H, Ugawa Y. Quadripulse stimulation (QPS). Exp Brain Res 2020;238:1619–25. https://doi.org/10.1007/s00221-020-05788-w.

[16] George MS, Lisanby SH, Avery D, Mcdonald WM, Durkalski V, Pavlicova M, et al. Daily Left Prefrontal Transcranial Magnetic Stimulation Therapy for Major Depressive Disorder. Arch Gen Psychiatry 2010;67:507–16.

[17] Thielscher A, Antunes A, Saturnino GB. Field modeling for transcranial magnetic stimulation: A useful tool to understand the physiological effects of TMS? Proc Annu Int Conf IEEE Eng Med Biol Soc EMBS 2015:222–5. https://doi.org/10.1109/EMBC.2015.7318340.

[18] Zmeykina E, Mittner M, Paulus W, Turi Z. Weak rTMS-induced electric fields produce neural entrainment in humans. Sci Rep 2020;10:1–16. https://doi.org/10.1038/s41598-020-68687-8.

[19] Beynel L, Davis SW, Crowell CA, Dannhauer M, Lim W, Palmer H, et al. Site-specific effects of online rtms during a working memory task in healthy older adults. Brain Sci 2020;10:1–19. https://doi.org/10.3390/brainsci10050255.

[20] Kraft A, Dyrholm M, Kehrer S, Kaufmann C, Bruening J, Kathmann N, et al. TMS over the right precuneus reduces the bilateral field advantage in visual short term memory capacity. Brain Stimul 2015;8:216–23. https://doi.org/10.1016/j.brs.2014.11.004.

[21] Liu A, Vöröslakos M, Kronberg G, Henin S, Krause MR, Huang Y, et al. Immediate neurophysiological effects of transcranial electrical stimulation. Nat Commun 2018. https://doi.org/10.1038/s41467-018-07233-7.

[22] Alekseichuk I, Mantell K, Shirinpour S, Opitz A. Comparative modeling of transcranial magnetic and electric stimulation in mouse, monkey, and human. Neuroimage 2019;194:136–48. https://doi.org/10.1016/j.neuroimage.2019.03.044.

[23] Shirinpour S, Hananeia N, Rosado J, Galanis C, Vlachos A, Jedlicka P, et al. Multi-scale Modeling Toolbox for Single Neuron and Subcellular Activity under (repetitive) Transcranial Magnetic Stimulation. BioRxiv 2020:2020.09.23.310219. https://doi.org/10.1101/2020.09.23.310219.

[24] Sykes M, Matheson NA, Brownjohn PW, Tang AD, Rodger J, Shemmeii JBH, et al. Differences in motor evoked potentials induced in rats by transcranial magnetic stimulation under two separate anesthetics: Implications for plasticity studies. Front Neural Circuits 2016;10. https://doi.org/10.3389/fncir.2016.00080.

[25] Bungert A, Antunes A, Espenhahn S, Thielscher A. Where does TMS Stimulate the Motor Cortex? Combining Electrophysiological Measurements and Realistic Field Estimates to Reveal the Affected Cortex Position. Cereb Cortex 2017;27:5083–94. https://doi.org/10.1093/cercor/bhw292.

[26] Laakso I, Murakami T, Hirata A, Ugawa Y. Where and what TMS activates : Experiments and modeling. Brain Stimul 2018;11:166–74. https://doi.org/10.1016/j.brs.2017.09.011.

[27] Wilson MT, George LS. Repetitive Transcranial Magnetic Stimulation : A Call for Better Data. Front Neural Circuits 2016;10:57. https://doi.org/10.3389/fncir.2016.00057.

[28] Siebner HR, Ziemann U. What is the threshold for developing and applying optimized procedures to determine the corticomotor threshold? Clin Neurophysiol 2014;125:1–2. https://doi.org/10.1016/j.clinph.2013.07.012.

[29] Gilio F, Rizzo V, Siebner HR, Rothwell JC. Effects on the right motor hand-area excitability produced by low-frequency rTMS over human contralateral homologous cortex. J Physiol 2003;551:563–73. https://doi.org/10.1113/jphysiol.2003.044313.

[30] Pridmore S, Fernandes Filho JA, Nahas Z, Liberatos C, George MS. Motor threshold in transcranial magnetic stimulation: A comparison of a neurophysiological method and a visualization of movement method. J ECT 1998;14:25–7. https://doi.org/10.1097/00124509-199803000-00004.

[31] Schutter DJLG, Honk JV. A Standardized Motor Threshold Estimation Procedure for Transcranial Magnetic Stimulation Research. J ECT 2006;22:176–8.

[32] Rounis E, Yarrow K, Rothwell JC. Effects of rTMS conditioning over the fronto-parietal network on motor versus visual attention. J Cogn Neurosci 2007;19:513–24. https://doi.org/10.1162/jocn.2007.19.3.513.

[33] Lang N, Harms J, Weyh T, Lemon RN, Paulus W, Rothwell JC, et al. Stimulus intensity and coil characteristics influence the efficacy of rTMS to suppress cortical excitability. Clin Neurophysiol 2006;117:2292–301. https://doi.org/10.1016/j.clinph.2006.05.030.

