## Supplemental for "Selecting stimulation intensity in repetitive transcranial magnetic stimulation studies: A systematic review between 1991 and 2020"

*Shared senior authorship.

**Correspondance**: zsolt.turi[at]anat.uni-freiburg.de

| Year | Total nr. of articles | Step 1 | Step 2 |
|---|---|---|---|
| 1991 | 1 | 1 | 0 |
| 1992 | 0 | 0 | 0 |
| 1993 | 3 | 3 | 0 |
| 1994 | 9 | 9 | 0 |
| 1995 | 2 | 2 | 0 |
| 1996 | 6 | 6 | 0 |
| 1997 | 6 | 6 | 0 |
| 1998 | 14 | 14 | 0 |
| 1999 | 27 | 14 | 13 |
| 2000 | 36 | 15 | 15 |
| 2001 | 56 | 15 | 15 |
| 2002 | 65 | 15 | 15 |
| 2003 | 70 | 15 | 15 |
| 2004 | 91 | 15 | 15 |
| 2005 | 98 | 15 | 15 |
| 2006 | 133 | 15 | 15 |
| 2007 | 137 | 15 | 15 |
| 2008 | 137 | 15 | 15 |
| 2009 | 150 | 15 | 15 |
| 2010 | 182 | 15 | 15 |
| 2011 | 217 | 15 | 20 |
| 2012 | 212 | 16 | 20 |
| 2013 | 256 | 16 | 20 |
| 2014 | 266 | 16 | 20 |
| 2015 | 260 | 16 | 20 |
| 2016 | 281 | 16 | 20 |
| 2017 | 263 | 16 | 20 |
| 2018 | 283 | 16 | 20 |
| 2019 | 298 | 16 | 20 |
| 2020 | 228 | 16 | 22 |
| **Sum** | **3,787** | **380** | **380** |

**Table S1. The number of processed articles per year in each sampling step.** Since the number of articles depends on the year of publication, we used year-dependent rule-governed sampling. In the first sampling step, we selected all articles between 1991 and 1998, because in these early years we identified only a few articles (n = 41). In 1999, we randomly took 14 articles and between 2000 and 2010, we randomly picked 15 articles in each year (n = 165), because there were less than 200 publications per year. Between 2011 and 2020, we randomly selected 16 articles (n = 160). In the second sampling step, we selected 13 articles for the year 1999, since we identified only 27 articles in this year. Moreover, we randomly picked 15 articles in each year between 2000 and 2010 (n = 165 articles) and 20 articles between 2011 and 2019 (n = 180) and 22 articles for year 2020.

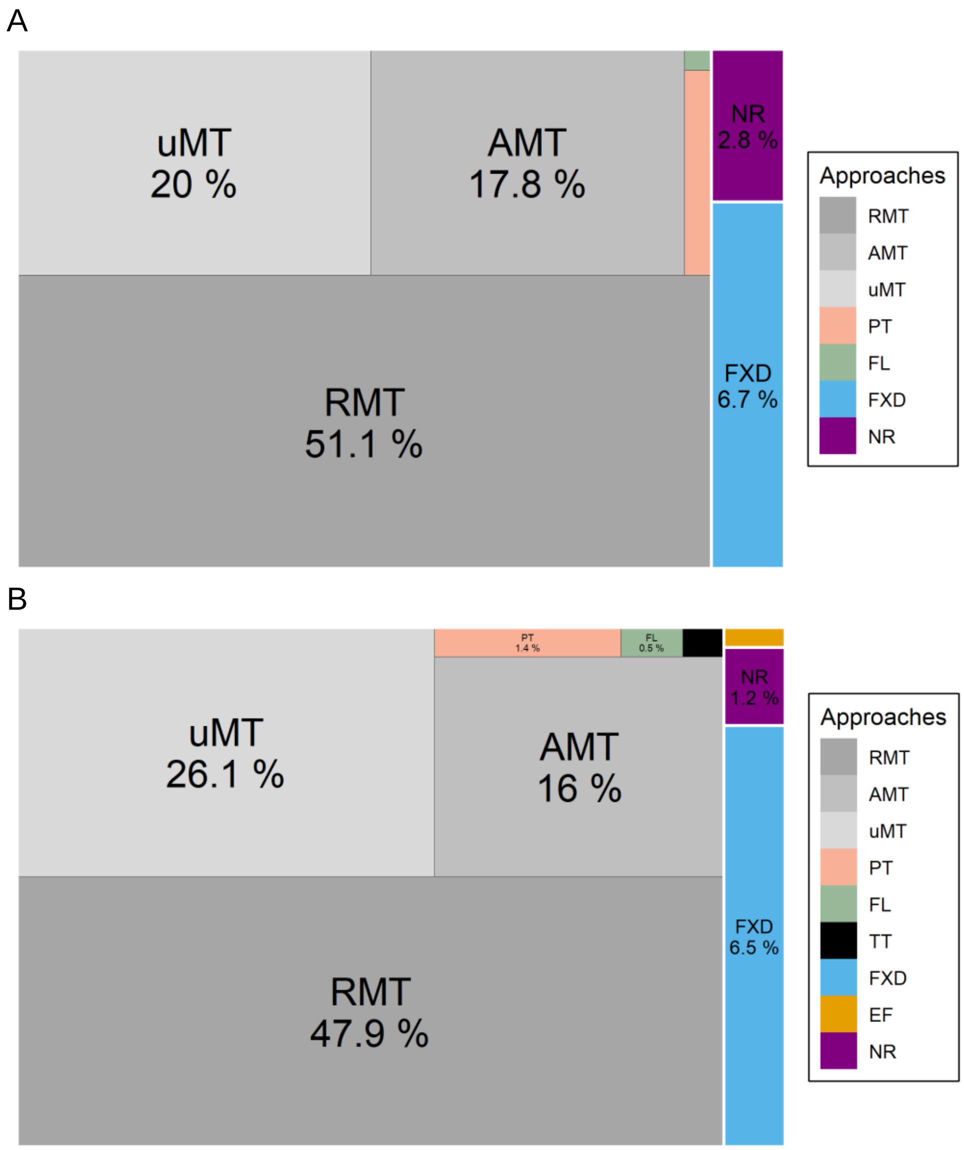


**Fig. S1: The stimulation intensity selection approaches and their relative frequency of use in the first sample (A) and in the second (B).** In each sample, we appraised 380 articles, respectively. Percentages calculated as relative the total number of identified protocols. The sizes of rectangles are proportional to the percentage values. Abbreviations: RMT: resting motor threshold, AMT: active motor threshold, uMT: unspecified motor threshold, PT: phosphene threshold, FL: functional lesion threshold, TT: tolerability threshold, FXD: fixed intensity, EF: electric-field based intensity selection, NR: not reported.

| Feature | Keywords |
|---|---|
| Motor threshold | “active motor threshold”, “resting motor threshold”, “rest motor threshold”, “motor threshold”, “AMT”, “RMT” and “MT” |
| Device output | "maximal stimulator output", "maximum stimulator output", "maximal power", "maximum power", "maximal output", "maximum output", "maximal capacity", "maximum capacity", "device capacity", "device's capacity", "machine capacity", "stimulator output", "machine output", "device output", " MSO", "fixed intensity" |
| Coil | "figure-8", "figure-of-8", "figure eight", "figure-eight", "figure-of-eight", "figure of eight", "eight-shaped", "eight shaped", "butterfly", "H-coil", "H coil", "H1-coil", "H1 coil", "double-cone", "double cone". |
| Stimulator | "Cadwell", "Brainsway", “CR Technology”, "Dantec", "Deymed", "eNeura", "EB Neuro", "Langer Medical", "Mag and More", "Mag & More", "Magstim", "Mag-Stim", "Magventure", "Medtronic", "Neuronetics", "Nanjing Weisi", "Neurosoft", "Neurostar", "Nexstim", “Nihon Kohden”, "Remed", “Yingchi Technology”, "Yiruide", "Yunsheng" |
| Electric field* | "V/m", "mV/mm", "electric field", "electric-field", "electrical field", "electrical-field", "e-field" and "E-field" |

**Table S2. The keywords used for the text-mining search. In all cases, the search result was manually processed**. *We performed the search for the electric field on the entire database. For the rest of the features, we limited the search on the 20% of the randomly selected articles.
